# Thrombospondin-4 mediates hyperglycemia- and TGF-beta-induced inflammation in breast cancer

**DOI:** 10.1101/2020.01.03.894436

**Authors:** Santoshi Muppala, Roy Xiao, Jasmine Gajeton, Irene Krukovets, Dmitriy Verbovetskiy, Olga Stenina-Adognravi

## Abstract

Inflammation drives the growth of tumors and is an important predictor of cancer aggressiveness. CD68, a marker of tumor-associated macrophages (TAM), is routinely used as a marker to aid in prognosis and treatment choices for breast cancer.

We report that thrombospondin-4 (TSP-4) mediates breast cancer inflammation and growth in mouse models in response to hyperglycemia and TGF-beta by increasing TAM infiltration and production of inflammatory signals in tumors. Analysis of breast cancers and non-cancerous tissue specimens from hyperglycemic patients revealed that levels of TSP-4 and of macrophage marker CD68 are upregulated in diabetic tissues. TSP-4 was co-localized with macrophages in cancer tissues. Bone-marrow-derived macrophages (BMDM) responded to high glucose and TGF-beta by upregulating TSP-4 production and expression, as well as the expression of inflammatory markers.

We report a novel function for TSP-4 in breast cancer: regulation of TAM infiltration and inflammation. The results of our study provide new insights into regulation of cancer growth by hyperglycemia and TGF-beta and suggest TSP-4 as a potential therapeutic target.

**Novelty and Impact:** Thrombospondin-4 (TSP-4) is a secreted extracellular protein that belongs to the family of matricellular proteins. TSP-4 is one of the top 1% of proteins upregulated in several cancers, including breast cancer. Inflammation and infiltration of macrophages drive cancer progression and metastasis and are clinically important markers of cancer aggressiveness and critical consideration in the process of selection of the appropriate therapeutic approaches. We report that TSP-4 promotes breast cancer inflammation and infiltration of macrophages and mediates the effects of hyperglycemia and TGF-beta on cancer growth and inflammation. Our work describes a role for TSP-4 in cancer inflammation and identifies the pathways, in which increased levels of TSP-4 mediate cancer growth.

## 1. Introduction

Cancer growth depends on the interplay between the cancer cells and the tumor microenvironment. Tumor inflammation is modulated by the infiltration of immune cells and is closely associated with the tumor aggressiveness and metastasis (1). Activated cancer and vascular cells produce chemoattractants and pro-inflammatory signals to recruit inflammatory cells from blood. The accumulation of inflammatory cells in a tumor is an important prognostic index that has been successfully used to evaluate the aggressiveness of cancer in conjunction with other indexes describing the proliferation rate of cancer cells and their migratory potential. For example, one of the 12 markers evaluated in Oncotype DX (a test used to predict the aggressiveness of breast cancer and to make therapeutic decisions) is CD68, a marker of macrophages (2).

The composition of the extracellular matrix (ECM) regulates recruitment of immune cells and their interactions with other cell types in a tumor. In recent years, the role of ECM, and of matricellular proteins specifically, in regulation of inflammatory cell functions attracted more attention (3–5). Thrombospondins (TSP), the matricellular proteins that are not structural but regulate cell-ECM and cell-cell interactions, belong to a family that consists of five distinct members with distinct properties and roles in inflammation (6). While the functions of TSP-1 in regulation of inflammation have been somewhat examined, the roles for other TSPs are unknown. TSP4 is especially interesting as a potential regulator of inflammation, due to its highly upregulated levels in several conditions linked to inflammation, including coronary artery disease, heart failure, and cancers (3,7–12). Indeed, we found that TSP-4 regulates cancer growth (13,14) and promotes pro-inflammatory polarization of macrophages (5), in addition to regulation of the blood and vascular cells adhesion, migration, and activation (3,5). However, the role of TSP-4 in regulation of cancer inflammation and infiltration of tumors with macrophages remained unknown. Here, we report that TSP-4 regulates cancer inflammation by increasing tumor infiltration with tumor-associated macrophages (TAM) and the levels of pro-inflammatory markers.

A rapidly-growing body of evidence connects hyperglycemia, insulin resistance, and glucose intolerance with cancer risk and progression (15–24). Diabetes, pre-diabetes, and metabolic syndrome are associated with chronic inflammation (25,26), and high glucose promotes pro-inflammatory polarization of macrophages (27,28) *in vitro*. To understand the role of TSP-4 as a mediator of hyperglycemia-induced cancer inflammation, we examined TSP-4 levels in breast cancer xenografts in mouse models of hyperglycemia and in the breast cancer tissues of hyperglycemic patients, as well as the effects of high glucose on cultured macrophages.

TGF-beta is a major regulator of tissue remodeling and inflammation and the main regulator of ECM production (29,30), a function critical in many pathological conditions, including cancer. As a cytokine with well-studied receptors and intracellular signaling, TGF-beta and the pathways it activates are attractive therapeutic targets. However, TGF-beta has cell- and process-specific effects that are sometimes distinct and even opposite, depending on a stage and localization of a process (so-called “TGF-beta paradox) (31). In cancer, TGF-beta suppresses the growth of tumor at the early stages by inducing apoptosis and cell cycle arrest in cancer cells and by supporting cell differentiation. However, at the later stages associated with increased inflammation and angiogenesis, TGF-β promotes tumor growth and metastasis by modulating the tumor microenvironment. We found that TGF-beta is a potent inducer of TSP-4 production in endothelial cells and macrophages (5,14). Thus, one of the goals of the current report was to demonstrate the role of TSP-4 as a process-specific mediator of TGF-beta effects on cancer inflammation.

Our report demonstrates the effect of TSP-4 on cancer inflammation and the role of TSP-4 as a mediator of the effects of hyperglycemia and TGF-beta, two recognized drivers of cancer inflammation, growth, and metastasis with incompletely understood mechanisms of action.

## 2. Materials and Methods

### 2.1. Patient’s breast cancer and adjacent normal tissue specimens

were obtained from the Cleveland Clinic Tissue Bank. The work was approved by the Cleveland Clinic Institutional Review Board. All patients were female. Patients with HbAc1 < 6 were considered normoglycemic; patients with HbAc1 > 7 or documented diabetes diagnosis were considered hyperglycemic. The patient information is summarized in Supplemental Tables 1 and 2. H&E stained sections of tumors from normoglycemic and hyperglycemic patients are shown in Suppl. Fig. 1 (top four panels).

### 2.2. Animals

Mice were of the C57BL/6 background. Both genders were used. *Thbs4^-/-^* and P387-TSP4-KI mice were described previously (5). Animal procedures were approved by the IACUC and in agreement with the NIH Guide for Animal Use.

### 2.3. Induction of hyperglycemia in mice

Hyperglycemia was induced by streptozotocin (STZ) injections (32,33). Mice with levels ≥250 mg/dl were included in the experiments.

### 2.4. EMT6 mouse breast cancer xenografts

Fifteen-to sixteen-week-old mice were anesthetized by IP injection of Ketamine (80mg/kg)/Xylazine (15mg/kg) mixture. Mice were injected in the mammary fat pad with 1.5 × 10^6^ EMT6 cells (RRID:CVCL_1923) (ATCC) in 100 μl of saline. All experiments were performed with mycoplasma-free cells. TGF-β1 (500ng/kg) injections were given IP daily for 14 days. The injected amount of TGF-β1 was based on reported studies in mice (34,35) and our previous work (14). Tumors were harvested at the end of the experiment. H&E stained representative sections of EMT6 xenografts from normoglycemic and hyperglycemic mice are shown in Suppl. Fig. 1, four bottom panels.

### 2.5. Bone Marrow Derived Macrophages (BMDM)

BMDM were isolated from tibia of WT and *Thbs4^-/-^* mice as described by others (36) and in our previous publication (5) and cultured in M-CSF (14-8983-62, ThermoFisher)/DMEM/F12 for 5 days. 95-98% of cultured cells were identified as macrophages by immunohistochemistry and FACS analysis.

BMDM were treated either with D-Glucose (30mM) or TGF-beta (10ng/ml) for 6h and 24h respectively, along with proper controls. The concentrations of D-Glucose and TGF-beta were selected based on previous studies in BMDM and other cultured cells (5,14,32,33,37–42). 30 mM D-Glucose was producing the maximal effects while still reflecting a naturally occurring blood level of glucose *in vivo*.

### 2.6. Quantitative real-time RT-PCR analysis

Total RNA was extracted using TRIzol Reagent (#15596026, ThermoFisher) followed by cDNA synthesis using SuperScript^™^ First-Strand Synthesis System (#11904018, ThermoFisher). Taqman fast Master mix (#4444557, ThermoFisher). The TaqMan^®^ probes for specific gene products, with *ActB, Gapdh*, and *Rn18s* as housekeeping gene controls.

### 2.7. Immunohistochemistry, immunofluorescence, confocal imaging and quantification of macrophage markers

10 μm sections of mouse tumors were stained with primary rat anti-mouse CD68 antibody (MCA1957B, Bio-Rad), and goat anti-human TSP-4 (AF2390, R&D Systems). Sections of human tumors were stained with mouse anti-human CD68 (Dako) and goat anti-human TSP-4 (AF2390, R&D Systems) using Vecta Stain ABC Kit (PK-6104, PK-6102, PK-6105 Vector). Visualization after staining with the antibodies was performed using a high-resolution slide scanner (Leica SCN400FL, Leica microsystems, GmbH, Wetzlar, Germany) at 20 × magnification. For immunofluorescence, Collagen I (Ab6308, Abcam) and Collagen IV (Ab6586, Abcam), rat anti-mouse CD68 (MCA1957B, Bio-Rad), goat anti-human TSP-4 (AF2390, R&D Systems), monoclonal mouse anti-human alpha smooth muscle actin (M0851, Dako), monoclonal mouse anti-human CD68 (M0814, Dako), rabbit anti-human vWF antibody (A0082, Dako), were used with corresponding secondary antibodies. Images were taken at a high-resolution confocal microscope (Leica DM 2500).

High-resolution images of the whole section were generated and quantified to determine the percentage of the stained area using ImagePro 6.1. Quantification was performed by investigator blinded to the assignment of animals between groups. The number of pixels in the whole section of tissue (excluding area without tissue and large gaps in tissue specimen) was defined as total area. The stained area (colored) was selected and quantified in pixels, and the percent of stained are was calculated.

Representative H&E images of tumors from mice and patients are shown in Suppl. Fig. 1.

### 2.8. Statistical analysis

Analyses of the data were performed using Sigma Plot Software (Systat Software, San Jose, CA, USA): Student’s t-test and ANOVA were used to determine the significance of parametric data, and Wilcoxon rank sum test was used for nonparametric data. The significance level was set at p=0.05. The data are presented as mean±S.E.M.

## 3. Results

### 3.1. TSP-4 deletion reduces inflammation in mouse breast cancer

To understand the role of TSP-4 in cancer inflammation, EMT6 cells transduced with TSP-4 shRNA (13) were injected into WT or *Thbs4^-/-^* mice (Fig. 1A). Although we could not detect TSP-4 production by cultured EMT6, we found that EMT6 cells do produce trace amounts of TSP-4 *in vivo*, and the best inhibition of BC tumor growth was observed with BC cells with suppressed TSP-4 production (13).

**Figure 1.**
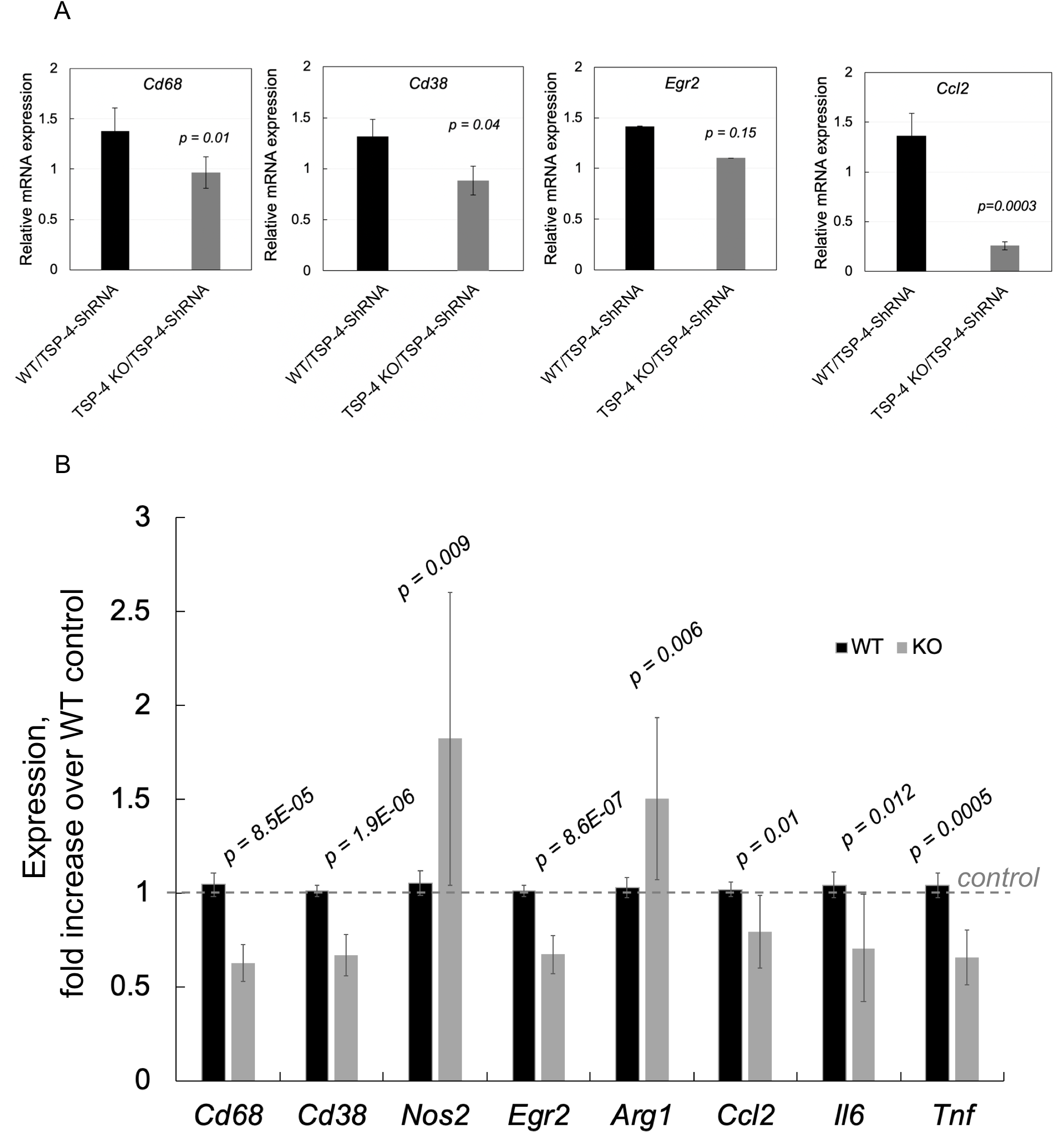
TSP-4 promotes macrophage infiltration and inflammation in vivo and in cultured bone marrow-derived macrophages (BMDM). **A:** *Cd68, Cd38, Egr2*, and *Ccl2* (MCP-1) mRNA levels were measured in EMT6 mouse breast cancers grown in WT and *Thbs4^-/-^* mice. Real Time RT-PCR, n = 10, mean± S.E.M. **B:** Expression of *Thbs4, Il6, Ccl2, Tnf*, macrophage marker *Cd68*, pro-inflammatory macrophage markers *Cd38* and *Nos2*, and tissue repair macrophage markers *Egr2* and *Arg1* was measured in EMT6 mouse breast cancers xenografts from WT and *Thbs4^-/-^* mice.

The expression of a macrophage marker *Cd68*, a marker of pro-inflammatory macrophages *Cd38*, a marker of tissue resolving macrophages *Egr2*, and *Ccl2* (MCP-1) was quantified by Real-Time RT-PCR (Fig. 1A). *Cd68* and pro-inflammatory markers *Cd38* and *Ccl2* expression was decreased in *Thbs4^-/-^* mice, indicating that TSP-4 promotes macrophage infiltration of tumors and inflammation. *Cd68* expression was decreased from 1.36±0.19 to 0.75±0.11, p = 0.01; *Ccl2* (MCP-1) expression was decreased from 1.37±0.22 to 0.26±0.03, p = 0.0003, indicating a lower level of inflammation in the absence of TSP-4.

### 3.2. Effect of TSP-4 deletion on expression of inflammatory markers in cultured bone-marrow-derived macrophages (BMDM)

Expression of markers of macrophage polarization and inflammatory markers was measured in BMDM from WT and *Thbs4^-/-^* mice (Fig. 1B). Deletion of *Thbs4* resulted in significantly decreased expression of most markers, with the exception of one of the markers of pro-inflammatory macrophages *Nos2* and one of the markers of tissue repair macrophages *Arg1*: both were significantly upregulated in BMDM from TSP-4 KO mice.

### 3.3. Increased TSP-4 expression in breast cancer tumors of hyperglycemic animals

We reported that hyperglycemia accelerates the growth and angiogenesis in mouse models [40]. To understand the role of TSP-4 in regulations of hyperglycemia-induced inflammation that also drives the cancer growth, two mouse models of hyperglycemia, STZ-treated WT C57BL/6 mice (model of type 1 diabetes) and *Lepr^db/db^* (genetic model of type 2 diabetes) were used, and TSP-4 expression and protein levels were examined in EMT6 mouse breast cancer xenografts (Fig. 2 A and B). RNA was purified from the tumors, and levels of mRNA of thrombospondin-4 (*Thbs4*) were measured by RealTime RT-PCR (Fig. 2A and B, top left panels, and Supplemental Table 3). TSP-4 expression was significantly upregulated in tumors of both mouse models.

**Figure 2.**
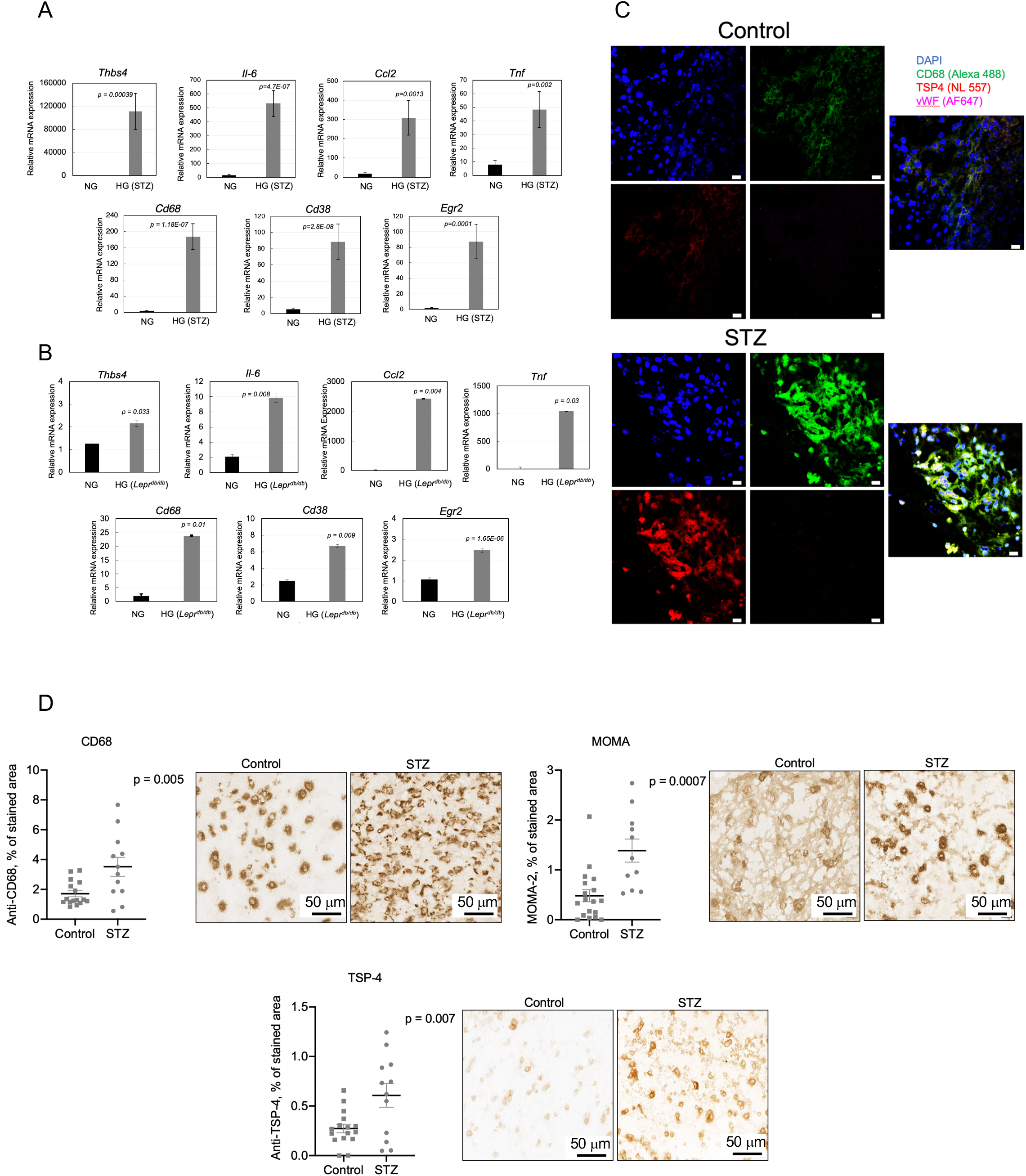
Increased levels of TSP-4, inflammatory markers and infiltration of macrophages in breast cancer tumors of hyperglycemic mice. EMT6 mouse breast cancer cells were injected into mammary fat pad of hyperglycemic WT STZ-treated **(A)** n=10, mean ± S.E.M and *Lepr^db/db^* **(B)** mice as described in methods, and mRNA levels of *Thbs4, Ccl2, Tnf*, macrophage marker *Cd68*, inflammatory macrophage marker *Cd38*, and tissue repair macrophage marker *Egr2* were measured by Real Time RT-PCR in tumors. NG = normoglycemic, HG = hyperglycemic; n=5, mean ± S.E.M.; **C:** Immunofluorescent staining of CD68, TSP-4 and vWF, (red: *Thbs4* stained with anti– TSP-4, green: macrophages stained with anti-CD68 antibody, magenta: vWF stained with anti-vWF; blue: nuclei stained with DAPI). Immunostaining of EMT6 tumors harvested from normoglycemic control (upper panel) and hyperglycemic (STZ-treated, bottom panel) WT C57BL/6 mice; **D:** TSP-4 and macrophages were stained with corresponding antibodies as described in Methods. Immunohistochemistry images and quantification of protein levels in tumors from hyperglycemic (STZ) and normoglycemic (Control) mice. n = 10, Mean ± S.E.M; scale bar = 20 μM.

EMT6 tumors from hyperglycemic mice were stained with anti-TSP-4 antibody to examine localization of TSP-4 in tumors and the levels of TSP-4 proteins. TSP-4 protein levels were also significantly upregulated in response to hyperglycemia (Fig. 2C and 2D). In hyperglycemic STZ-treated mice, immunofluorescence detected increased levels of TSP-4 (Fig. 2C, red), and co-staining with anti-CD68 antibody against macrophage marker (green) revealed that TSP-4 was co-localized with CD68, a marker of macrophages.

### 3.3. Increased TSP-4 expression and protein levels in breast cancer tumors of hyperglycemic patients

Immunohistochemistry was used to detect and quantify the levels of TSP-4 protein in patients’ breast cancer tumors and non-cancerous adjacent breast tissue (Fig. 3). TSP-4 protein levels were higher in breast cancer tissue as compared to adjacent non-cancerous tissue and increased in both the cancerous and non-cancerous tissues of diabetic patients (Fig. 3A, right panel) (7.01±0.38 % of total area of a section in tumor vs 3.32±0.41 % in non-cancerous in hyperglycemic patient, p = 0.02 and 4.73±0.21 % in tumor vs 1.03±0.20 % in non-cancerous in normoglycemic patients, p = 0.03).

**Figure 3.**
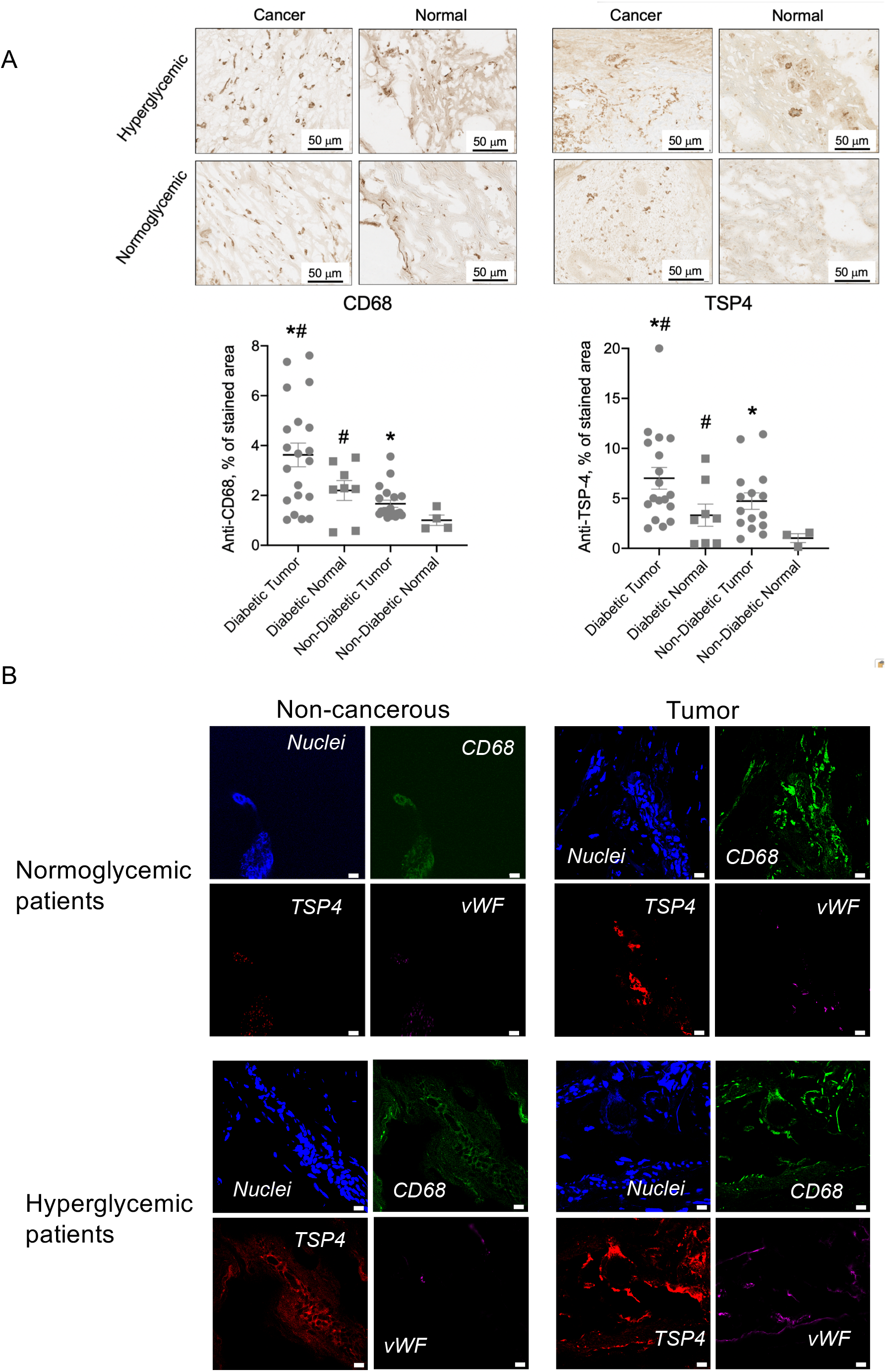
TSP-4 and macrophage marker CD68 in human diabetic tissues. **A:** TSP-4 and CD68 were stained with corresponding antibodies as described in Methods. Immunohistochemistry images and quantification of protein levels in tumors (Cancer) and adjacent non-cancerous tissue (Normal) from diabetic (Hyperglycemic) and nondiabetic (Hyperglycemic) patients. **B:** Immunoflurescence of tumors and adjacent non-cancerous tissues from normoglycemic (HbAc1<6) and hyperglycemic (HbAc1>7) patients; staining of CD68, TSP-4 and vWF, (red: *Thbs4* stained with anti–TSP-4, green: macrophages stained with anti-CD68 antibody, magenta: vWF stained with anti-vWF; blue: nuclei stained with DAPI).. Scale bar = 10 μM.

When breast cancer tissue specimens from hyperglycemic (HbAc1>7) and normoglycemic (HbAc1<6) patients were examined, most TSP-4 was associated with macrophages (Fig. 3B), although co-staining with a marker of endothelial cells (EC) vWF (purple) was also detected.

### 3.5. Increased TAM infiltration in breast cancer tumors of hyperglycemic animals and patients

mRNA of a macrophage marker CD68 was upregulated in EMT6 tumors of STZ-treated and *Lepr^db/db^* mice (Fig. 2A and B) suggesting increased number of macrophages in cancer xenografts in hyperglycemic mice.

Increased number of macrophages in tumors of hyperglycemic animals was detected by immunofluorescence and immunohistochemistry with anti-CD68 antibody (Fig. 2C and 2D): the area stained with anti-CD68 increased from 1.70±0.19 % of total area of a section in control normoglycemic mice to 3.51±0.62 % in STZ-treated hyperglycemic mice, p = 0.005 (Fig. 2D, top left panel). Immunohistochemistry with MOMA-2, another antibody recognizing mouse macrophages, supported the data obtained with anti-CD68 antibody: area stained with MOMA-2 antibody significantly increased in tumors from hyperglycemic mice (Fig. 2D, top right panel).

CD68 protein levels were higher in breast cancer tissue of patients as compared to adjacent non-cancerous tissue (stained area 3.63±0.20 % of total section area in tumor vs 2.2±0.27 % in non-cancerous in hyperglycemic patient, p = 0.04 and 2.26±0.10 % in tumor vs 1.01±0.20 % in non-cancerous in normoglycemic patients, p = 0.01) (Fig. 3A, left panels). The levels of CD68 were significantly higher in diabetic tissues, in both the cancerous and the non-cancerous adjacent tissue (Fig. 3A, left panels)..

In both the animal and patients’ tumors, TSP-4 was associated predominantly with macrophages (Fig. 2C and 3B).

### 3.4. Increased inflammation in EMT6 mouse breast cancer of hyperglycemic mice

The levels of all inflammatory and TAM markers were significantly increased in tumors from STZ-treated mice (Fig. 2A - D and Supplemental Table 3), indicating a higher level of inflammation and increased TAM infiltration in tumors of hyperglycemic animals but also suggesting that tissue repair macrophages have been recruited into tumors along with pro-inflammatory macrophages.

All inflammation and macrophage markers were upregulated in hyperglycemic *Lepr^db/db^* mice (Fig. 2B and Supplemental Table 3).

### 3.6. Effect of TSP-4 deletion on the expression of inflammatory markers by BMDM in response to high glucose

The role of TSP-4 in high-glucose-induced upregulation of inflammatory markers was tested using BMDM isolated from WT mice and from *Thbs4^-/-^* mice (Fig. 4A and Suppl. Table 4). Deletion of TSP-4 prevented the high-glucose-induced increases in mRNA levels of macrophage marker *Cd68*, of markers of pro-inflammatory macrophages *Cd38* and *Nos2*, a marker of tissue-repair macrophages *Arg1*, and increase in *Tnf* levels without an effect on *Egr2, Ccl2*, and *Il6*.

**Figure 4.**
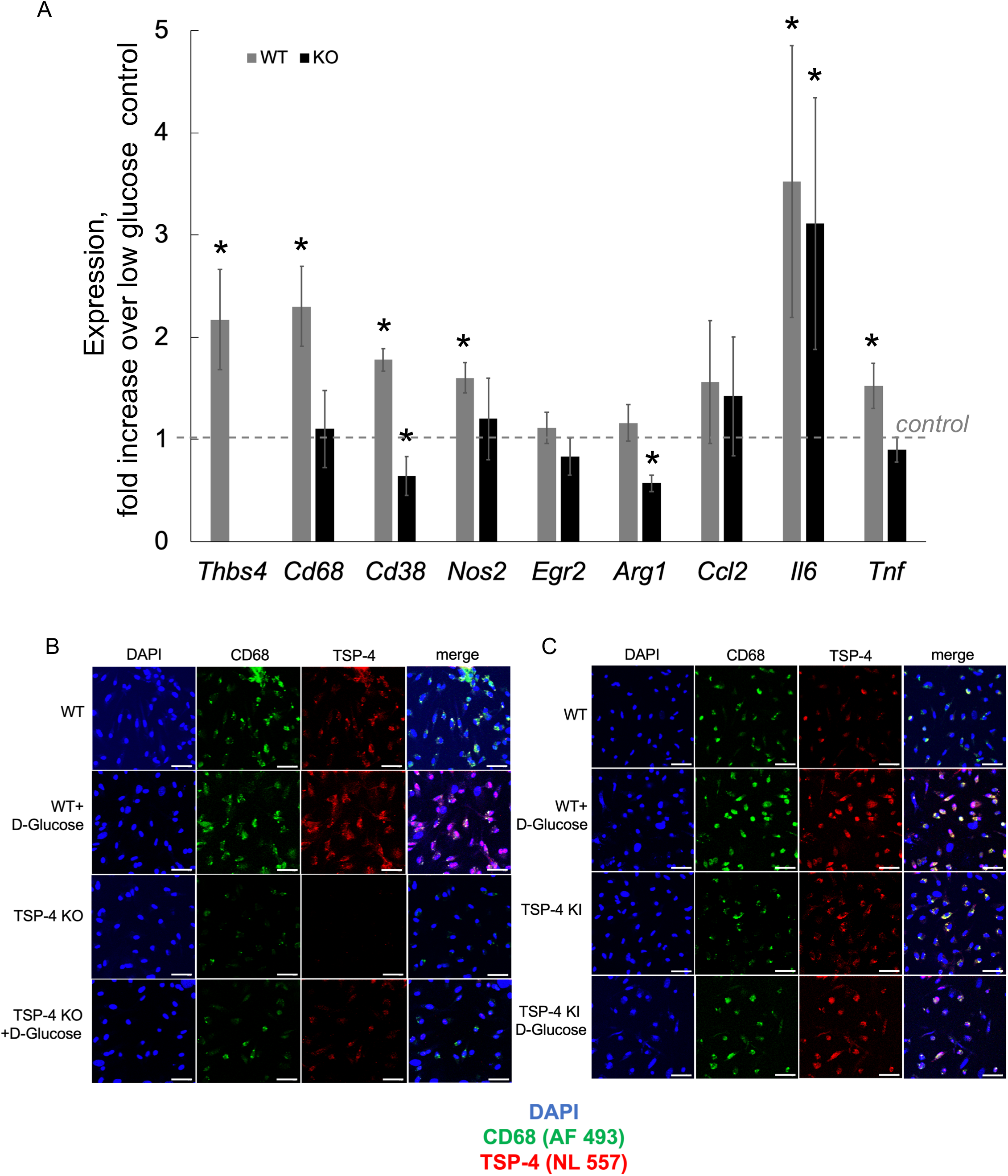
TSP-4 mediates the effects of high glucose on the levels of inflammatory markers and TSP-4 in cultured BMDM. Cultured BMDM from WT and *Thbs4^-/-^* (TSP-4 KO) mice were stimulated with 25 mM D-glucose for 24 h. **A:** mRNA levels of *Thbs4, Ccl2, Tnf, Il6*, macrophage marker *Cd68*, inflammatory macrophage markers *Cd38 and Nos2*, and tissue repair macrophage markers *Egr2* and *Arg1* were measured by Real Time RT-PCR; n=3; mean± S.E.M; *p<0.05 in comparison to control of the same genotype. **B, C:** TSP-4 and macrophage marker CD68 were visualized in cultured BMDM using corresponding antibodies. BMDM from WT mice, *Thbs4^-/-^* mice (TSP-4 KO, **B**) and mice expressing more active form of TSP-4, P387 TSP-4 (TSP-4 KI, **C**) were stimulated with 25 mM L- or D-glucose for 24 h. Blue: nuclei, DAPI; green: anti-Cd68; red: anti-TSP-4. Scale bar = 30 μM.

### 3.7. Effect of high glucose on TSP-4 protein production in cultured bone-marrow-derived macrophages (BMDM)

Because most of TSP-4 protein was co-localized with TAM in tumors, we isolated bonemarrow-derived macrophages (BMDM) from WT, *Thbs4^-/-^* (TSP-4 KO mice), and P387-TSP-4 KI mice expressing a mutant form of TSP-4 that is more active in all cellular interactions (4,5,13,14,40,43,44). Cultured BMDM were incubated in low (5 mM) D-glucose or high (30 mM) D-glucose and stained with anti-CD68 and anti-TSP-4 (Fig. 4B and C).

TSP-4 levels were increased in BMDM isolated from WT mice in response to high glucose and were higher in cultured BMDM from P387-TSP-4-KI mice, consistent with our previous observations of higher stability and activity of the mutant protein (43,44). High glucose did not increase the levels of CD68 in P387-TSP-4-KI mice. The deficiency in TSP-4 (*Thbs4^-/-^*, TSP-4 KO) reduced the level of CD68 (Fig. 4B).

### 3.8. TGF-beta increases inflammation in mouse breast cancer

Previously, we reported that TGF-beta accelerates the growth of EMT6 xenografts and cancer angiogenesis in mice, and that TSP-4 is a mediator of TGF-beta effects on cancer angiogenesis [14]. Because cancer inflammation drives the tumor growth, we examined the effect of TGF-beta1 injections on TSP-4 and inflammation in mouse EMT6 cancer. Mice with EMT6 xenografts were injected intraperitoneally with 500 ng/kg TGF-beta daily. The expression of TSP-4 gene and inflammation markers was examined by Real-Time RT-PCR (Fig. 5A and Supplemental Table 5). Daily TGF-beta1 injections resulted in increased levels of *Thbs4, Cd68, Il6, Ccl2* (MCP-1), and *Tnf*. The levels of a marker of pro-inflammatory macrophages *Cd38* were increased by TGF-beta1 injections, while the level of a marker of tissue-repair macrophages *Egr1* was decreased.

**Figure 5.**
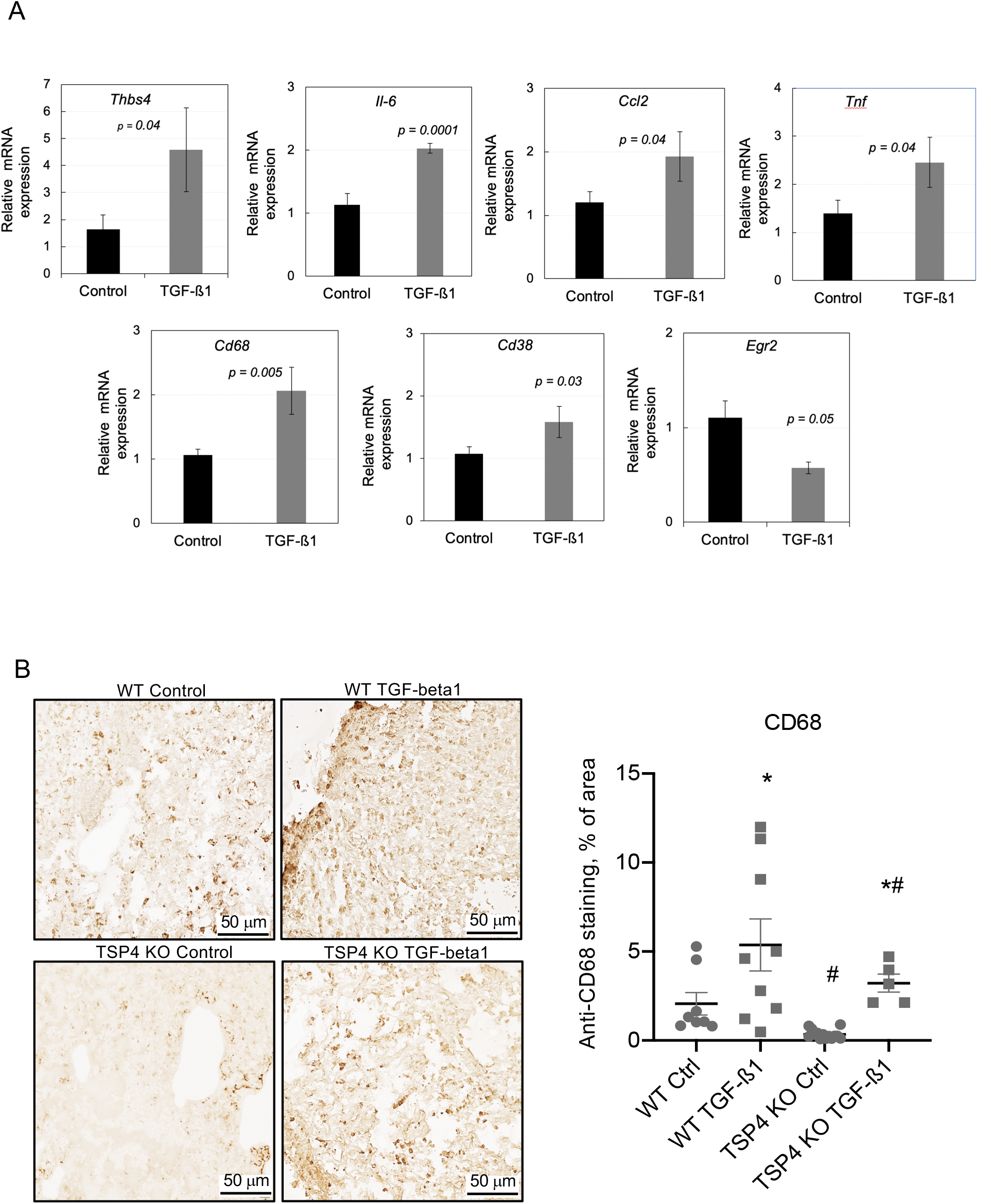
TGF-beta1 increases inflammation in mouse breast cancer. **A:** Expression of *Thbs4, Il6, Ccl2, Tnf*, a macrophage marker *Cd68*, a pro-inflammatory macrophage marker *Cd38*, and a tissue repair macrophage marker *Egr2* was measured in EMT6 mouse breast cancers of mice injected with TGF-beta as described in Methods. n = 10, Mean ± S.E.M. **B:** CD68 was visualized in EMT6 tumors of WT and *Thbs4^-/-^* mice injected with TGF-beta1, and the stained area was quantified as described in Methods. Immunohistochemistry; n = 10, mean ± S.E.M; *p<0.05 as compared to corresponding no treatment control, #p<0.05 as compared to the same treatment in different genotype.

### 3.9. Increased macrophage infiltration in breast cancer tumors of mice injected with TGF-beta1

Increased number of macrophages in tumors of animals injected with TGF-beta1 was detected by immunohistochemistry with anti-CD68 antibody (Fig. 5B). The area stained with anti-CD68 antibody was decreased in *Thbs4^-/-^* mice.

### 3.10. Effect of TSP-4 deletion on the expression of inflammatory markers by BMDM in response to TGF-beta1

Effect of TSP-4 KO was examined in cultured BMDM stimulated with TGF-beta1 (Fig. 6A and Suppl. Table 6). The expression of *Cd68* and *Cd38* was only modesty upregulated by TGF-beta1 in WT cells, and this modest upregulation was prevented by TSP-4 deletion. Nos2 levels were significantly increased by TGF-beta1 in WT cells, and the increase was completely prevented in BMDM from *Thbs4^-/-^* mice. Increase in *Arg1, Il6* and *Tnf* expression was significant and unaffected by the lack of TSP-4. *Egr1* and *Ccl2* expression was decreased by TGF-beta1 in cells from mice of both genotypes.

**Figure 6.**
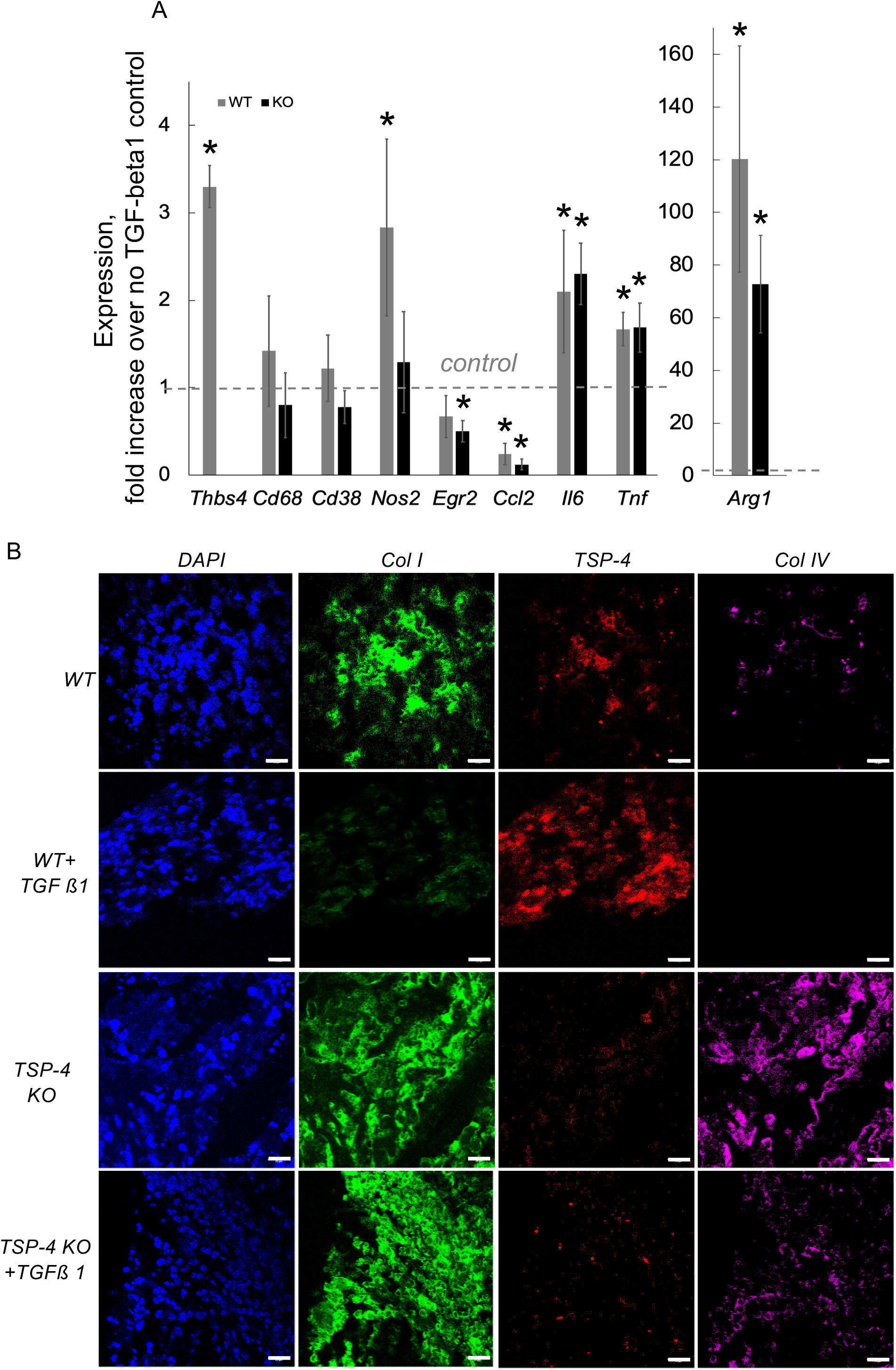
**A: Effects of TGF-beta1 on the levels of inflammatory markers in cultured BMDM.** Cultured BMDM from WT and *Thbs4^-/-^* (TSP-4 KO) mice were stimulated with 10ng/ml of TGF-beta1 for 24 h, n=10; mean± S.E.M; *p<0.05 in comparison to control of the same genotype; **B: Effect of TSP-4 KO on collagen production in breast cancer in response to TGF-beta1.** TSP-4 (red), nuclei (blue), collagen I (green) and collagen IV (magenta) were visualized using corresponding antibodies. Immunofluorescence, scale bar = 50 μM.

### 3.11. TSP-4 does not mediate the effects of TGF-beta on collagen production

To investigate whether TSP-4 mediates other effects of TGF-beta1 in breast cancer tumors, in addition to regulation of angiogenesis and inflammation, we examined collagen I and IV (Col I and IV) levels in tumors grown in WT and *Thbs4^-/-^* mice (Fig. 6B). Both Col I and Col IV levels were increased in *Thbs4^-/-^* mice, consistent with our previous observation of suppression of collagen production by TSP-4 (45). TGF-beta1 decreased the level of Col I in a TSP-4-dependent manner: the decrease was completely prevented in *Thbs4^-/-^* mice. The levels of Col IV were decreased in response to TGF-beta1 injections independently of the presence or the absence of TSP-4, suggesting that TSP-4 does not mediate the effect of TGF-beta on collagen IV production or turnover.

## 4. Discussion

TSP-4 is one of the top upregulated genes in several cancers, including the breast cancer (12,46,47). Using *Thbs4^-/-^* mice, we demonstrated that TSP-4 deletion prevents the growth of cancer and mediates TGF-beta1 effect on cancer angiogenesis (14). Tumor inflammation is an important marker of cancer aggressiveness and is defined by the tumor microenvironment, including the composition of the tumor ECM. We recently reported that TSP-4 promotes the accumulation of macrophages in tissues and regulates their pro-inflammatory polarization, adhesion, and migration (3,5). Taking in the account the dramatically increased levels of TSP-4 in some cancers, its effects on tumor growth, and its role in regulation of macrophage polarization and function, we investigated here whether TSP-4 regulates cancer inflammation.

Two well-known regulators of ECM production, inflammation, and cancer growth - hyperglycemia (15,16,48) and TGF-beta (14,49,50) - are known to regulate the remodeling of tissues and of cancer microenvironment. Both have poorly understood effects on cancer growth and ECM. We have examined these two stimuli to define the role of TSP-4 in pro-inflammatory effects of TGF-beta and hyperglycemia and found that in both pathways TSP-4 mediates macrophage infiltration and cancer inflammation and growth.

First, we documented that TSP-4 promotes infiltration of macrophages in EMT6 tumors and inflammation. The deletion of TSP-4 in mice (*Thbs4^-/-^*) and EMT6 cancer cells [EMT6 stably transduced with TSP-4 shRNA] resulted in significantly reduced expression of macrophage marker CD68, pro-inflammatory macrophage marker CD38, CCl2 (MCP-1), but not a marker of tissue repair macrophages Egr2, consistent with regulation of macrophage infiltration and upregulation of inflammation by TSP-4 (3,5).

Experiments in cultured BMDM supported pro-inflammatory effect of TSP-4: the expression of *Ccl2, Il6, Tnf*, and *Cd38* was reduced in BMDM from TSP-4 KO mice. However, these experiments suggested a more complex effect of TSP-4 on macrophage differentiation, because the levels of the general marker of macrophages *Cd68* was also reduced, and another marker of pro-inflammatory macrophages, *Nos2*, was upregulated in TSP-4 KO cells. The complexity of the effects on differentiation was also clear from the inconsistent effects on both the markers of pro-inflammatory macrophages (*Cd38* levels were decreased while *Nos2* levels were increased in BMDM from the TSP-4 KO mice) and tissue repair macrophages (*Egr2* levels were decreased while *Arg1* levels were increased in BMDM from the TSP-4 KO mice). The differential effects on the specific markers suggest that TSP-4 promotes a very specific pathway of polarization of macrophages favoring a subtype that is not the major pro-inflammatory subtype. The complexity of TSP-4 role in macrophage polarization and differential roles of TSP-4 in pathways initiated by various inflammatory stimuli was discussed in our previous publication (5).

Several markers of inflammation were examined in breast cancer tissues of hyperglycemic mice and patients: inflammatory cytokines IL-6, CCL2 (MCP-1) and TNF-alpha, as well as a macrophage marker CD68, markers of pro-inflammatory macrophages CD38 and iNOS, and markers of tissue repair anti-inflammatory macrophages EGR2 and ARG1. Whether hyperglycemia was the result of streptozotocin (STZ) treatment or a genetic mutation in the leptin receptor (type 1 and type 2 diabetes models, respectively), similar increases in TSP-4 levels, infiltration of macrophages, and increased levels of all inflammatory markers were detected, documenting higher cancer inflammation in tumors of hyperglycemic mice. In our previous work, we also found that high glucose *in vitro* and hyperglycemia *in vivo*, independently of a used model, drives the cancer growth (5,32,33).

Similarly, higher macrophage infiltration of tumors and adjacent non-cancerous tissues was observed in specimens of hyperglycemic patients, parallel to the increased levels of TSP-4 in these specimens. We have previously reported that TSP-4 can be produced by endothelial, smooth muscle cells, and by macrophages (3,5,11,13,44). Most of TSP-4 in tumors was associated with macrophages, and only a small portion was colocalized with endothelial cells, suggesting that macrophages are the main or at least the major producers of TSP-4 in tumors.

Using BMDM isolated from WT mice, *Thbs4^-/-^* mice, and mice expressing a more active form of the protein P387-TSP-4 (*P387-TSP4-KI*), we tested the effect of high glucose on the levels of TSP-4. High glucose increased the production of TSP-4 by BMDM in WT cells.

To prove the role of TSP-4 as a mediator of proinflammatory effects of high glucose and TGF-beta, cultured BMDM from WT and *Thbs4^-/-^* mice were stimulated with high glucose or with TGF-beta1. Although the deletion of TSP-4 clearly resulted in lower production of inflammatory markers by the cultured BMDM, the effects were distinct and specific, suggesting that the effects of high glucose are not mediated by TGF-beta but rather that the two stimuli activate non-overlapping pro-inflammatory pathways. Remarkably, the effect on the expression of Ccl2 (MCP-1) in BMDM was opposite in cells stimulated with high glucose and TGF-beta: high glucose increased it, but TGF-beta1 decreased the expression.

*In vivo* in EMT6 xenografts, TGF-beta1 injections increased macrophage infiltration as was detected by the staining of the tumor sections with anti-CD68 antibody. The macrophage infiltration was decreased in tumors of *Thbs4^-/-^* mice, demonstrating that TSP-4 mediates the cancer inflammation.

Although both hyperglycemia and TGF-beta stimulations *in vivo* and in cultured BMDM resulted in increased expression and protein levels of TSP-4, the differential effects of the two stimuli on cultured BMDM and differential roles of TSP-4 in these effects of high glucose and TGF-beta1 suggested that glucose effects TSP-4 levels in TGF-beta-independent manner. The roles of TSP-4 in responses of BMDM to TGF-beta1 and high glucose were different: while both stimuli upregulated the expression of TSP-4, TSP-4 mediated the upregulation of *Ccl2* in response to TGF-beta1 with no effect of TSP-4 KO on upregulation *Il6* and *Tnf. Tnf* upregulation was down in TSP-4 KO cells in response to high glucose, but there was no effect of TSP-4 deletion on *Ccl2* and *Il6* upregulation by glucose. High glucose caused upregulation of markers of pro-inflammatory macrophages without affecting markers of tissue repair macrophages, and upregulation of all macrophage markers in response to glucose was reduced in TSP-4 KO cells. TGF-beta1 produced a more complex response, with *Nos2* and *Arg1* being highly upregulated with no effects on *Cd38* and reduction in *Egr2* expression, but TSP-4 KO resulted in reduction of the levels of all macrophage markers. ========

Stiffening of the tumor ECM due to increased levels of collagen and fibronectin is an important event in cancer progression. Although TSP-4 mediates TGF-beta1 effects on cancer growth, angiogenesis, and inflammation, TSP-4 involvement is process-specific. When we examined the effects of TSP-4 deletion on regulation of collagen depositions in response to TGF-beta1, we found that levels of collagen IV are regulated in TSP-4-independent manner: TGF-beta1 downregulated collagen IV depositions in both the WT and *Thbs4^-/-^* mice. TSP-4 deletion prevented the effect on collagen I levels.

Our work describes a role for TSP-4 in cancer inflammation and identifies the pathways, in which increased levels of TSP-4 in tumor may promote cancer growth.

## Abbreviations

*ActB*: beta-actin gene
ANOVA: Analysis of variance
Arg1: Arginase 1
BMDM: bone-marrow-derived macrophages
Ccl2: chemokine ligand 2, also known as monocyte chemoattractant protein 1
CD38: Cluster of Differentiation 38
CD68: Cluster of Differentiation 68
cDNA: complementary DNA (deoxyribonucleic acid)
Col I: collagen I
Col IV: collagen IV
DMEM: Dulbecco's Modified Eagle Medium
EC: endothelial cell
ECM: extracellular matrix
Egr2: early growth response protein 2
*Gapdh*: Glyceraldehyde 3-phosphate dehydrogenase gene
HbAc1: glycated hemoglobin
Il-6: interleukin-6
iNOS: Inducible Nitric Oxide Synthase
IP: intraperitoneal
M-CSF: Macrophage Colony Stimulating Factor
MCP-1: macrophage chemoattractant protein
mRNA: Messenger RNA
NIH: National Institutes of Health
Nos2: Nitric Oxide Synthase 2 (Inducible Nitric Oxide Synthase) gene
P387-TSP4-KI: knock-in mice with A to P mutation in position 387
*Rn18s*: 18S ribosomal RNA
RT-PCR: Reverse transcription polymerase chain reaction
S.E.M.: standard error of the mean
STZ: streptozotocin
TAM: tumor-associated macrophages
TGF-beta: Transforming Growth Factor-beta
*Thbs4*: TSP-4 gene
TNF-alpha: Tumor necrosis factor alpha
TSP-4: thrombospondin-4
vWF: von Willebrand factor
WT: wild type

## Acknowledgments

This work was supported by the National Institutes of Health awards R01 HL117216 and R01 CA177771 and by the American Heart Association award 17PRE33660475.

## Conflict of Interests

The authors received funding for the described work. This work was supported by the National Institutes of Health awards R01 HL117216 and R01 CA177771 to Olga Stenina-Adognravi and by the American Heart Association award 17PRE33660475 to Jasmine Gajeton (PI) and Olga Stenina-Adognravi (sponsor).

There are no other conflicts to disclose.

## Ethical statement

The work with patients’ specimens was approved by the Cleveland Clinic Institutional Review Board. Animal procedures were approved by the IACUC and were in agreement with the NIH Guide for Animal Use.

## Data availability

The data that support the findings of this study are available from the corresponding author upon reasonable request.

---

**Suppl. Fig. 1.**
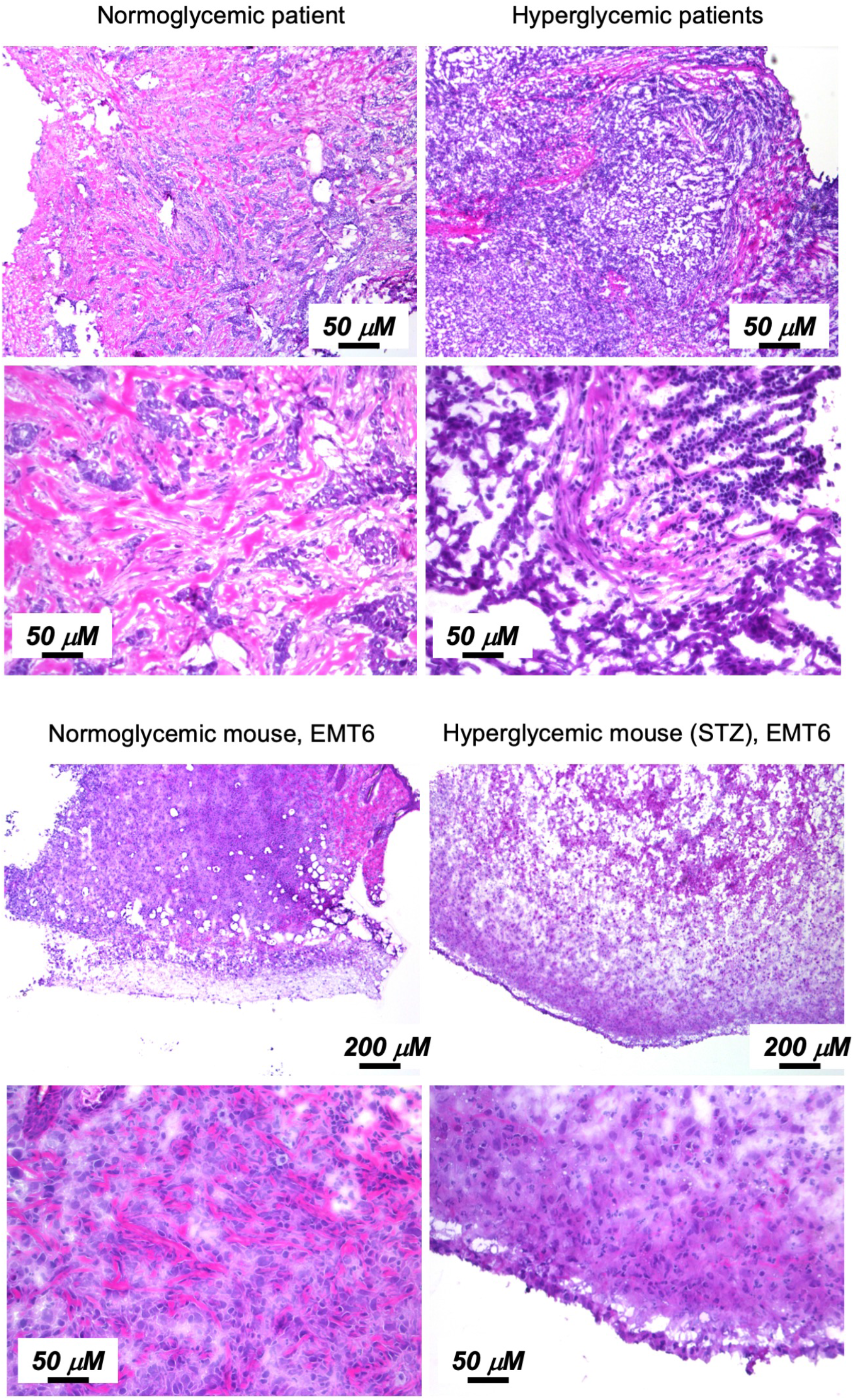
H&E stained sections of tumors from normoglycemic and hyperglycemic patients and mice.

**Supplemental Table 1.**
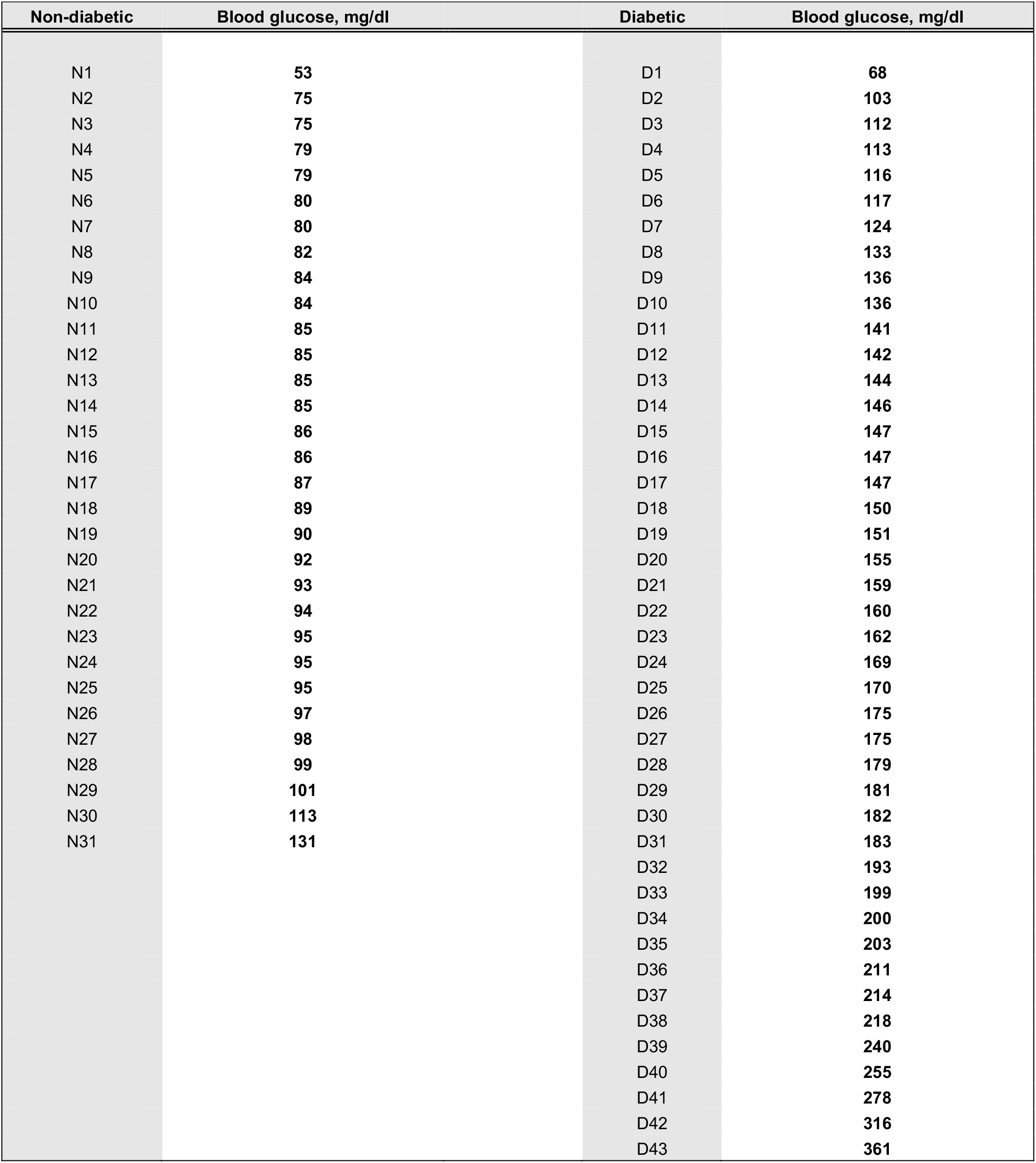
Blood glucose levels in non-diabetic (N1 – N31) and diabetic (D1 – D43) breast cancer patients, mg/dl.

| **Non-diabetic** | **Blood glucose, mg/dl** | **Diabetic** | **Blood glucose, mg/dl** |
| --- | --- | --- | --- |
| N1 | 53 | D1 | 68 |
| N2 | 75 | D2 | 103 |
| N3 | 75 | D3 | 112 |
| N4 | 79 | D4 | 113 |
| N5 | 79 | D5 | 116 |
| N6 | 80 | D6 | 117 |
| N7 | 80 | D7 | 124 |
| N8 | 82 | D8 | 133 |
| N9 | 84 | D9 | 136 |
| N10 | 84 | D10 | 136 |
| N11 | 85 | D11 | 141 |
| N12 | 85 | D12 | 142 |
| N13 | 85 | D13 | 144 |
| N14 | 85 | D14 | 146 |
| N15 | 86 | D15 | 147 |
| N16 | 86 | D16 | 147 |
| N17 | 87 | D17 | 147 |
| N18 | 89 | D18 | 150 |
| N19 | 90 | D19 | 151 |
| N20 | 92 | D20 | 155 |
| N21 | 93 | D21 | 159 |
| N22 | 94 | D22 | 160 |
| N23 | 95 | D23 | 162 |
| N24 | 95 | D24 | 169 |
| N25 | 95 | D25 | 170 |
| N26 | 97 | D26 | 175 |
| N27 | 98 | D27 | 175 |
| N28 | 99 | D28 | 179 |
| N29 | 101 | D29 | 181 |
| N30 | 113 | D30 | 182 |
| N31 | 131 | D31 | 183 |
|  |  | D32 | 193 |
|  |  | D33 | 199 |
|  |  | D34 | 200 |
|  |  | D35 | 203 |
|  |  | D36 | 211 |
|  |  | D37 | 214 |
|  |  | D38 | 218 |
|  |  | D39 | 240 |
|  |  | D40 | 255 |
|  |  | D41 | 278 |
|  |  | D42 | 316 |
|  |  | D43 | 361 |

**Supplemental Table 2.**
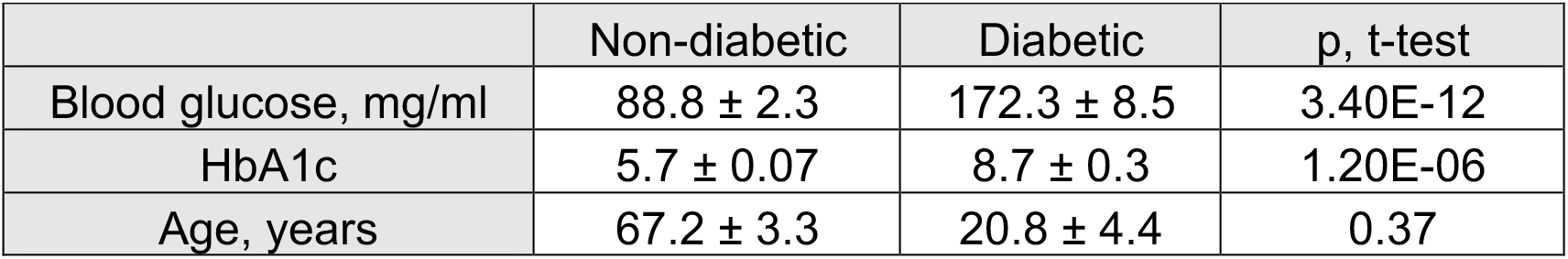
Blood glucose levels, HbA1c levels, and age of patients. Mean ± S.E.M.; non-diabetic, n=31; diabetic, n = 43.

|  | Non-diabetic | Diabetic | p, t-test |
| --- | --- | --- | --- |
| Blood glucose, mg/ml | 88.8 ± 2.3 | 172.3 ± 8.5 | 3.40E-12 |
| HbA1c | 5.7 ± 0.07 | 8.7 ± 0.3 | 1.20E-06 |
| Age, years | 67.2 ± 3.3 | 20.8 ± 4.4 | 0.37 |

**Supplemental Table 3.** Expression of TSP-4, macrophage markers, and inflammatory molecules in hyperglycemic mice. (Mean ± S.E.M.; N = normoglycemic, H = hyperglycemic).

| Model<br>Gene | STZ-treated mice |  |  | *Lepr<sup>db/db</sup>* mice |  |  |
| --- | --- | --- | --- | --- | --- | --- |
|  | N | H | P, t-test | N | H | P, ttest |
| *Thbs4* | 6.9±2.98 | 111124.9±31134.7 | 0.00039 | 1.26±0.16 | 2.15±0.35 | 0.033 |
| *Il6* | 16±6.2 | 532±93.2 | 4.7E-07 | 2.09±0.2 | 9.89±0.6 | 0.008 |
| *Ccl2* | 18.61±7.57 | 310±90 | 0.0013 | 4.13±1.4 | 2429.8±862.6 | 0.0044 |
| *Tnf* | 8.12±2.9 | 48.5±13.2 | 0.002 | 2.55±0.73 | 1046.1±544.8 | 0.032 |
| *Cd68* | 3.31±1.24 | 187.77±32 | 1.18E-07 | 2.06±0.52 | 23.84±8.86 | 0.01 |
| *Cd38* | 5.28±1.8 | 88.64±21.7 | 2.8E-08 | 2.50±0.63 | 6.73±0.87 | 0.009 |
| *Egr2* | 1.79±0.51 | 87.42±22.1 | 0.0001 | 1.07±0.09 | 2.47±0.22 | 1.65E-06 |

**Supplemental Table 4.** Expression of TSP-4, macrophage markers, and inflammatory molecules in BMDM in response to high glucose. (Mean ± S.E.M.; N = normoglycemic, H = hyperglycemic).

| Model<br>Gene | WT mice |  |  | *Thbs4<sup>-/-</sup>* mice |  |  |
| --- | --- | --- | --- | --- | --- | --- |
|  | Control | High glucose | P, t-test | Control | High glucose | P, ttest |
| *Thbs4* | 1.02±0.14 | 2.54±0.58 | 0.03 | N/A | N/A | N/A |
| *Il6* | 1±0.07 | 2.68±0.4 | 0.007 | 1.24±0.5 | 1.51±0.42 | 0.34 |
| *Ccl2* | 1.02±0.15 | 3.42±0.47 | 0.004 | 0.52±0.02 | 1.6±0.33 | 0.002 |
| *Tnf* | 6.9±2.98 | 111124.9±31134.7 | 0.37 | 0.95±0.1 | 0.68±0.16 | 0.11 |
| *Cd68* | 1.05±0.23 | 2.61±0.29 | 0.006 | 0.55±0.07 | 0.44±0.14 | 0.26 |
| *Cd38* | 1.01±0.11 | 1.92±0.13 | 0.003 | 0.36±0.033 | 0.11±0.02 | 0.001 |
| *Nos2* | 1.07±0.26 | 1.78±0.1 | 0.03 | 0.7±0.03 | 0.94±0.16 | 0.096 |
| *Egr2* | 1.01±0.1 | 1.8±0.3 | 0.03 | 0.66±0.07 | 0.42±0.065 | 0.03 |
| *Arg1* | 1.03±0.3 | 1.98±0.71 | 0.05 | 1.3±0.43 | 1.21±.29 | 0.44 |

**Supplemental Table 5.** Expression of TSP-4, macrophage markers, and inflammatory molecules in EMT6 xenografts in response to TGF-beta1. (Mean ± S.E.M.; Control = PBS injections; TGF-beta1 = TGF-beta1 injections).

| Model<br>Gene | WT mice |  |  |
| --- | --- | --- | --- |
|  | Control | TGF-beta1 | P, t-test |
| *Thbs4* | 1.62±0.54 | 5.21±1.53 | 0.04 |
| *Il6* | 1.13±0.17 | 2.02±0.07 | 0.00013 |
| *Ccl2* | 1.21±0.16 | 1.93±0.39 | 0.049 |
| *Tnf* | 1.39±0.26 | 2.45±0.52 | 0.040 |
| *Cd68* | 1.07±0.08 | 2.06±0.36 | 0.005 |
| *Cd38* | 1.08±0.10 | 1.58±0.24 | 0.035 |
| *Egr2* | 1.10±0.17 | 0.57±0.06 | 0.05 |

**Supplemental Table 6.** Expression of TSP-4, macrophage markers, and inflammatory molecules in BMDM in response to TGF-beta1. (Mean ± S.E.M.; Control and TGF-beta1).

| Model<br>Gene | WT mice |  |  | *Thbs4*<sup>-/-</sup> mice |  |  |
| --- | --- | --- | --- | --- | --- | --- |
|  | Control | TGF-beta1 | P, t-test | Control | TGF-beta1 | P, ttest |
| *Thbs4* | 1.02±0.13 | 3.37±0.25 | 0.0006 | N/A | N/A | N/A |
| *Il6* | 1.01±0.13 | 3.35±0.8 | 0.02 | 0.59±0.24 | 2.74±0.65 | 0.01 |
| *Ccl2* | 1.01±0.09 | 0.24±0.05 | 0.0009 | 1.34±0.07 | 0.17±0.02 | 4E-05 |
| *Tnf* | 1.04±0.2 | 1.87±0.2 | 0.02 | 0.7±0.2 | 1.38±0.4 | 0.09 |
| *Cd68* | 1.01±0.11 | 4.22±1.01 | 0.016 | 0.67±0.1 | 3.45±0.82 | 0.01 |
| *Cd38* | 1.00±0.07 | 1.28±0.16 | 0.09 | 0.94±0.07 | 0.7±0.16 | 0.11 |
| *NOS2* | 1.00±0.06 | 3.46±0.83 | 0.02 | 0.43±0.13 | 1.1±0.15 | 0.01 |
| *Egr2* | 1.00±0.06 | 0.68±0.034 | 0.003 | 0.77±0.05 | 0.63±0.08 | 0.009 |
| *Agr1* | 1.05±0.23 | 126.6±3.48 | 1.6E-06 | 1.00±0.07 | 55.63±8.9 | 0.002 |

## Notes

### Competing Interest Statement

The authors have declared no competing interest.

### Summary of Updates

New data added and revisions to the text.

